# Feedforward and feedback mechanisms cooperatively regulate rapid experience-dependent response adaptation in a single thermosensory neuron type

**DOI:** 10.1101/2023.12.05.570166

**Authors:** Tyler J. Hill, Piali Sengupta

## Abstract

Sensory adaptation allows neurons to adjust their sensitivity and responses based on recent experience. The mechanisms that mediate continuous adaptation to stimulus history over seconds to hours long timescales, and whether these mechanisms can operate within a single sensory neuron type, are unclear. The single pair of AFD thermosensory neurons in *C. elegans* exhibits experience-dependent plasticity in their temperature response thresholds on both minutes- and hours-long timescales upon a temperature upshift. While long-term response adaptation requires changes in gene expression in AFD, the mechanisms driving rapid response plasticity are unknown. Here, we show that rapid thermosensory response adaptation in AFD is mediated via cGMP and calcium-dependent feedforward and feedback mechanisms operating at the level of primary thermotransduction. We find that either of two thermosensor receptor guanylyl cyclases (rGCs) alone is sufficient to drive rapid adaptation, but that each rGC drives adaptation at different rates. rGC-driven adaptation is mediated in part via phosphorylation of their intracellular domains, and calcium-dependent feedback regulation of basal cGMP levels via a neuronal calcium sensor protein. In turn, cGMP levels feedforward via cGMP-dependent protein kinases to phosphorylate a specific subunit of the cGMP-gated thermotransduction channel to further regulate rapid adaptation. Our results identify multiple molecular pathways that act in AFD to ensure rapid adaptation to a temperature change, and indicate that the deployment of both transcriptional and non-transcriptional mechanisms within a single sensory neuron type can contribute to continuous sensory adaptation.

**Significance statement:** The nervous system must continuously adapt to the sensory environment in order to adjust response sensitivity. Although both short- and long-term response adaptation has been reported to occur within sensory neurons themselves, how temporally distinct plasticity mechanisms are coordinated within single sensory neurons is unclear. We previously showed that long-term adaptation of temperature responses in the single AFD thermosensory neuron pair in *C. elegans* is mediated via gene expression changes in this neuron type. Here we show that multiple second messenger-driven feedforward and feedback mechanisms act to drive rapid thermosensory adaptation in AFD. Our results indicate that modulation of thermotransduction molecules via both transcriptional and non-transcriptional mechanisms contribute to distinct temporal phases of adaptation in a single sensory neuron type.

## INTRODUCTION

Animals live in highly dynamic environments and are thus subjected to constantly varying sensory stimuli. A key feature of sensory responses is their ability to adapt to the prevailing stimulus intensity, thereby preventing response saturation and enabling continued detection of salient stimulus changes. Since stimuli can vary over time, it is particularly critical that sensory adaptation also operate over a range of timescales to allow for effective information processing (1, 2). Rapid sensory adaptation operating within seconds to minutes can act at the level of sensory responses themselves, whereas long-term behavioral adaptation acting over minutes to hours is typically mediated via rescaling input-output functions at the circuit level (eg. (3–8). However, sensory neuron responses have also been reported to exhibit both rapid- and slow sensory adaptation (eg. (5, 9–11), raising the question of how these adaptation mechanisms are coordinated within individual sensory neurons to alter their working range.

Molecular mechanisms of sensory adaptation have been extensively studied in visual and olfactory neurons. Intracellular calcium flux plays a critical role in mediating rapid sensory adaptation in both systems, although calcium-independent pathways have also been described (12–14). Recent work suggests that in addition to these mechanisms that function over milliseconds to seconds, the experience-dependent transcriptional state of olfactory neurons contributes to sensory adaptation over a period of hours to days (9, 15). Whether similar or distinct molecules are targeted within individual olfactory neuron types to drive these temporally distinct forms of adaptation remains to be fully described.

Sensory responses have also been shown to adapt on timescales from minutes to hours in single sensory neuron types in *C. elegans* (4, 10, 11, 16–19). The bilateral pair of AFD thermosensory neurons exhibits particularly complex and temporally well-defined modes of adaptation. The response threshold of this neuron type adapts as a function of the animal’s temperature experience on both fast minutes-long and slow hours-long timescales, and these neurons are responsive to temperature changes only above their adapted threshold (10, 17–20). Experience-dependent adaptation of both the thermosensory response and synaptic output thresholds of AFD allows animals to retain the ability to respond to small temperature changes and to navigate thermal gradients efficiently over a 10°C temperature range (21, 22). Thus, studying thermosensory adaptation in AFD provides an opportunity to dissect the molecular pathways that mediate adaptation on different timescales within a single sensory neuron type, and to relate these adaptation mechanisms to physiologically relevant behavioral outputs.

As in other sensory neurons, thermotransduction in AFD is mediated via cyclic nucleotide and calcium signaling (21, 22) (Figure 1A). Warming temperatures above the adapted temperature threshold activates a trio of rGCs, increases intracellular cGMP levels and opens cyclic nucleotide-gated (CNG) channels to permit calcium influx and neuronal depolarization (17, 23–26). This response is terminated via hydrolysis of cGMP by multiple PDEs (26, 27) (Figure 1A). We recently showed that long-term adaptation to a new warm growth temperature is mediated in part via calcium-dependent expression changes in multiple genes in AFD including the thermosensor rGCs (10, 28, 29). Although intracellular calcium levels have also been suggested to mediate rapid adaptation in AFD (17), the molecular mechanisms that drive adaptation on a minutes-long timescale in this neuron are unknown.

**Figure 1.**
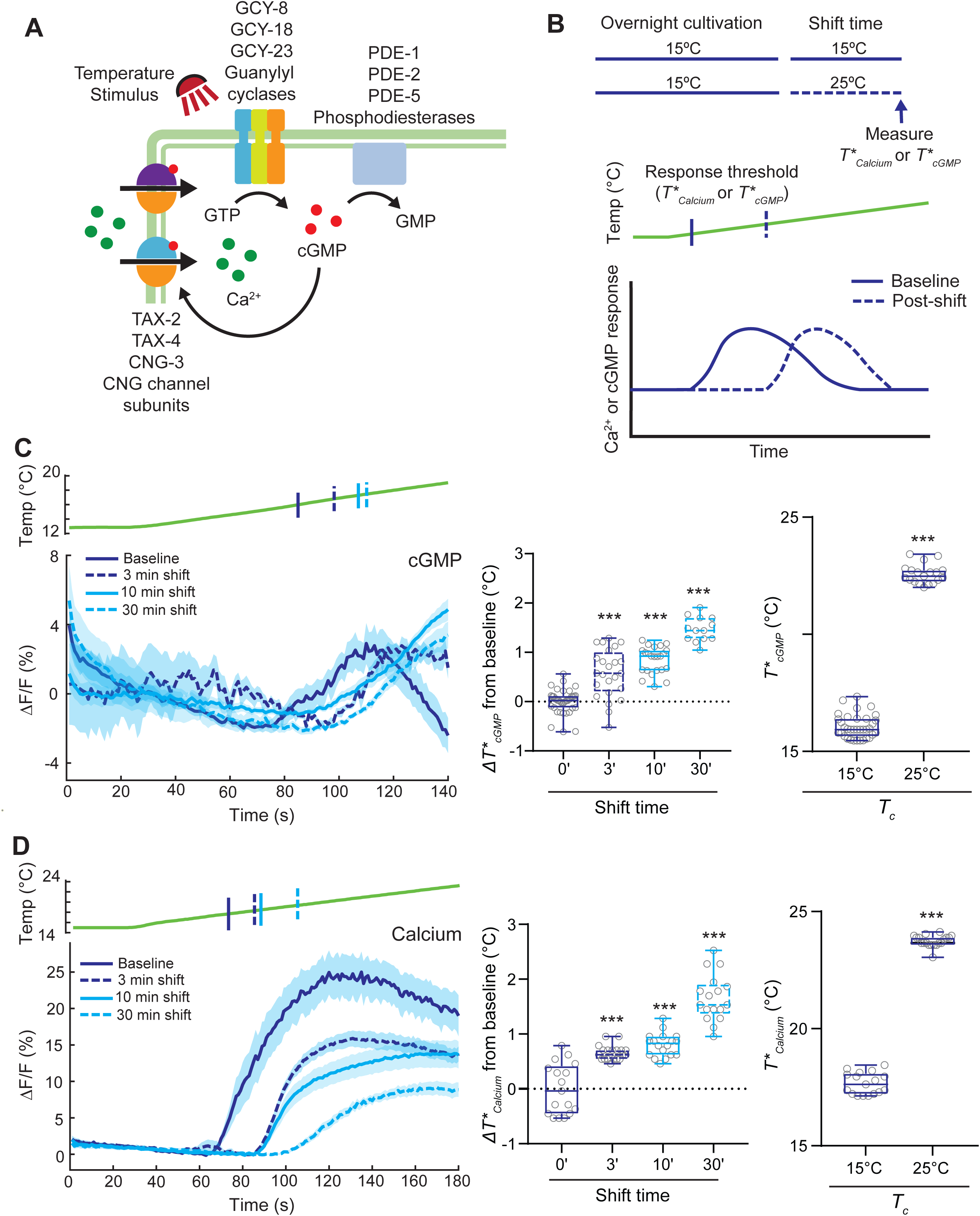
The thresholds of both intracellular cGMP and calcium response dynamics exhibit rapid temperature experience-dependent adaptation in AFD. **A)** Cartoon of the known thermotransduction pathway in AFD. Activation of the AFD-specific thermosensory receptor guanylyl cyclases (GCY-8, GCY-18, GCY-23) and/or inhibition of multiple phosphodiesterases (PDE-1 PDE-2, PDE-5) increases intracellular cGMP at temperatures above the adapted threshold. Opening of cGMP-gated channels (TAX-2, TAX-4, CNG-3) results in calcium influx and depolarization. See text for additional details and references. **B)** Cartoon of the temperature shift protocol. (Top) Animals grown overnight at 15°C are maintained at 15°C (baseline/no shift; solid line) or shifted to 25°C for different time periods (dashed line) prior to imaging. (Bottom) Schematic of calcium or cGMP response dynamics in AFD with no shift (solid line) or post temperature shift (dashed line) in response to a rising temperature ramp (green). *T*_cGMP_* and *T*_Calcium_*: threshold of the cGMP and calcium responses, respectively, pre- or post-temperature shift. **C,D)** (Left) Average changes in FlincG3 (C) or GCaMP6s (D) fluorescence in response to a rising temperature ramp (green; 0.05°C/sec linear ramp) in animals subjected to the shown temperature shift conditions. Shaded regions are SEM. Vertical lines on the temperature trace indicate the quantified average *T*_cGMP_* (C) or *T*_Calcium_* (D) (see Materials and Methods). (Middle) Average change in *T*_cGMP_* (C) or *T*_Calcium_* (D) from the baseline *T\** upon a temperature upshift for the indicated time periods. Each circle is the value from a single AFD neuron. Both cGMP and calcium measurements were performed at the AFD sensory endings. (Right) *T*_cGMP_* (C) or *T*_Calcium_* (D) of animals grown overnight at 15°C or 25°C (cultivation temperature, *T_c_*). Each circle is the value in a single neuron. n=15-30 neurons each; 2 independent experiments. Horizontal lines in boxes: median, lower and upper quartiles. Whiskers: maximum and minimum values. ***: different at p<0.001 from the 0’ (middle) or 15°C condition (right) (middle – ANOVA with Bonferroni correction; right – t-test).

Here we show that multiple calcium and cGMP-dependent mechanisms are integral to rapid response adaptation in AFD. We find that similar to the previously reported rapid adaptation of the calcium response, the threshold of cGMP responses also exhibits rapid response adaptation in AFD. While each of the examined thermosensor rGCs alone is sufficient to drive adaptation, each rGC mediates adaptation with different dynamics. Rapid adaptation is regulated via phosphorylation of the rGC intracellular domain, as well as regulation of basal cGMP levels; basal cGMP levels are modulated in part via calcium feedback via the NCS-2 neuronal calcium sensor protein. cGMP in turn feeds forward via the EGL-4 and PKG-2 cGMP-dependent protein kinases to phosphorylate the CNG-3 subunit of the CNG thermostransduction channels and further modulates the calcium response threshold in AFD. Together, our observations indicate that cGMP- and calcium-dependent feedforward and feedback mechanisms act in concert to mediate rapid experience-dependent plasticity in AFD thermosensory responses, thereby enabling this neuron to precisely adjust its response properties as a function of the animal’s temperature history.

## RESULTS

### The temperature-evoked cGMP response threshold in AFD exhibits rapid experience-dependent adaptation

Experience-dependent adaptation of the response threshold in AFD has been largely assessed via determining the temperature above which these neurons exhibit changes in intracellular calcium dynamics (*T*_Calcium_*) in response to a rising temperature ramp (10, 18–20) (Figure 1B). We considered two mutually non-exclusive mechanisms by which the threshold of temperature-evoked calcium influx via CNG channels in AFD is rapidly altered by the animal’s temperature history. In the first, the threshold of activation or inhibition of one or more of the AFD-expressed thermosensor rGCs (23, 24) or PDEs (27) (Figure 1A), respectively, may be reset by temperature history resulting in a shift in the gating threshold of the CNG channels. In the second, only the threshold of CNG channel opening may be reset. The first but not the second model predicts that similar to the adaptation of *T*_Calcium_*, the cGMP response threshold (*T*_cGMP_)* will also adapt rapidly upon a temperature change.

To distinguish between these models, we shifted adult animals from 15°C to 25°C for different periods of time, and compared *T*_cGMP_* and *T*_Calcium_* in AFD neurons expressing the genetically encoded FlincG3 or GCaMP6s cGMP and calcium sensors, respectively (18, 19, 25) (Figure 1B). Both *T*_cGMP_* and *T*_Calcium_* adapted to a significantly higher temperature within 3 mins of the temperature upshift from 15°C to 25°C (Figure 1C-D, Figure S1A-B), and continued to shift upwards upon further exposure to 25°C for up to 30 mins (Figure 1C-D, Figure S1A-B). As reported previously (18, 19, 25, 26), both *T*_cGMP_* and *T*_Calcium_* adapted to their final values upon overnight growth at 25°C (Figure 1C-D, Figure S1A-B). *T*_Calcium_* adaptation was similar at the AFD sensory endings and soma indicating that rapid adaptation occurs at the sensory endings themselves (Figure S1C). These observations indicate that similar to adaptation of *T*_Calcium_*, temperature experience also rapidly modulates *T*_cGMP_* in AFD. *T*_Calcium_* adaptation may be a consequence of *T*_cGMP_* adaptation alone, and/or due to modulation of CNG channels.

### Individual rGCs contribute differentially to rapid thermosensory adaptation in AFD

Temperature-regulated cGMP levels in AFD are determined by the opposing actions of cGMP production and hydrolysis by the GCY-8, GCY-18 and GCY-23 thermosensor rGCs, and multiple PDEs, respectively (Figure 1A) (23, 24, 27). All three rGCs contribute to correct adaptation of *T*_cGMP_* and *T*_Calcium_* following an overnight temperature upshift (23, 26, 30), but the roles of individual rGCs in regulating rapid thermosensory adaptation in AFD is unknown. Since misexpression of GCY-18 or GCY-23, but not GCY-8, is sufficient to confer temperature responses onto other cell types (23), we focused on the contributions of GCY-18 and GCY-23 to rapid thermosensory adaptation.

We examined cGMP and calcium responses in AFD neurons in *gcy-8 gcy-18* double mutants expressing GCY-23 alone, (Figure 2A-F), or in *gcy-23 gcy-8* double mutants expressing GCY-18 alone (Figure 2G-L). cGMP response amplitudes were significantly dampened in both double mutants as compared to responses in wild-type animals (Figure 2A,2G). Expression of either GCY-23 or GCY-18 alone was sufficient for rapid *T*_cGMP_* adaptation to warmer temperatures (Figure 2B,2H). However, *T*_cGMP_* in AFD expressing GCY-18 alone adapted to warmer temperatures more quickly than wild-type AFD following a temperature upshift (Figure 2H). *T*_cGMP_* was significantly lower in GCY-23- but not GCY-18-expressing neurons upon overnight growth at 25°C (Figure 2C,2I) (26). Consistent with lower levels of intracellular cGMP leading to the opening of fewer calcium channels, the amplitude of temperature-evoked calcium responses was also lower in neurons expressing either rGC (Figure 2D,2J). While *T*_Calcium_* also adapted rapidly upon a temperature upshift in neurons expressing either rGC alone, *T*_Calcium_* values were consistently lower and higher in GCY-23- and GCY-18-expressing neurons, respectively (Figure 2E,2K). *T*_Calcium_* was also lower in GCY-23- but not GCY-18-expressing animals upon overnight growth at 25°C (Figure 2F,2L). We previously showed that while misexpression of either rGC in chemosensory neurons is sufficient to confer temperature-evoked calcium responses, the threshold of activation of these responses in misexpressing neurons is not regulated by the long-term growth temperature (23). *T*_Calcium_* in ASE chemosensory neurons misexpressing GCY-23 also did not exhibit rapid response adaptation upon a temperature upshift (Figure S2), suggesting that the mechanisms driving this adaptation are likely to be regulated by AFD-specific pathways. We infer that expression of either rGC alone is sufficient to mediate rapid adaptation of AFD responses, but that each rGC drives adaptation at distinct rates.

**Figure 2.**
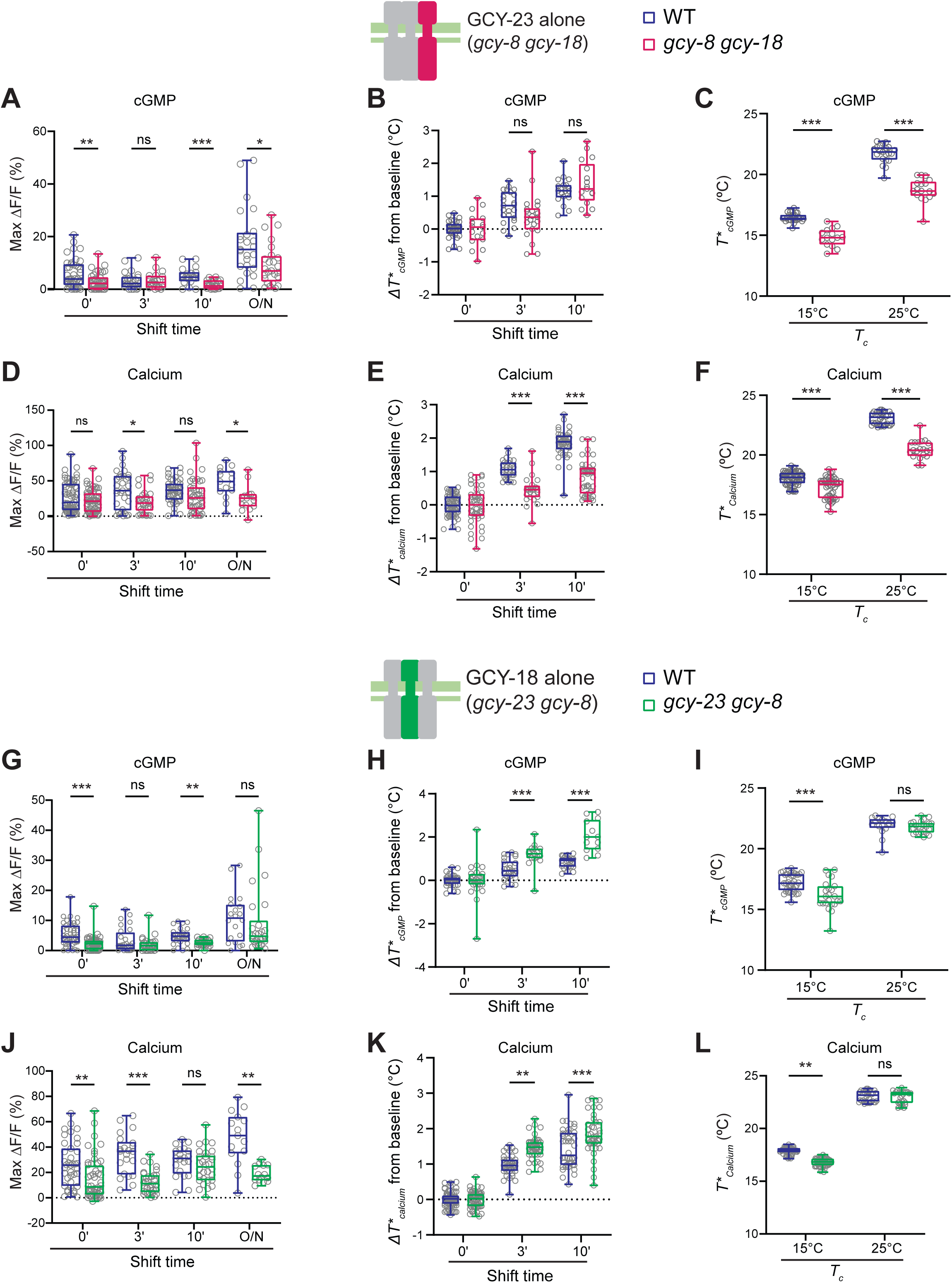
GCY-18 and GCY-23 drive thermosensory adaptation at distinct rates in AFD. **A,D,G,J)** Maximum FlincG3 (A,G) and GCaMP6s (D,J) response amplitudes in AFD in wild-type, *gcy-8 gcy-18* (expressing GCY-23 alone; A,D) or *gcy-23 gcy-8* (expressing GCY-18 alone; G,J) double mutant animals. Each circle is the amplitude of a single AFD neuron subjected to a rising temperature ramp under the indicated temperature exposure conditions. **B,E,H,K)** Average changes in *T*_cGMP_* (B,H) and *T*_Calcium_* (E,K) in response to the rising temperature ramps subjected to the shown temperature shift conditions in wild-type or double mutant animals. Each circle is the value in a single AFD neuron. **C,F,I,L)** *T*_cGMP_* (C,I) and *T*_Calcium_* (F,L) of wild-type and double mutant animals grown overnight at 15°C or 25°C (*T_c_*). Each circle is the value in a single animal. Horizontal lines in boxes in all panels: median, lower and upper quartiles. Whiskers: maximum and minimum values. *, **, and ***: different at p<0.05, 0.01, and 0.001, respectively between the indicated values (t-test). ns – not significant.

### The phosphorylation state of the rGC intracellular domain may modulate rapid adaptation and desensitization of their response threshold

How might a temperature shift reset the response threshold of a thermosensory rGC? The guanylyl cyclase A (GC-A) natriuretic peptide receptor is phosphorylated at six conserved residues in its intracellular juxtamembrane kinase homology domain in the unliganded state (31, 32). Ligand binding results in dephosphorylation of these residues thereby leading to response desensitization (33–36). Similar basal phosphorylation and inactivation via dephosphorylation upon ligand binding has also been reported for the sea urchin sperm guanylyl cyclase chemoreceptor (37, 38). We hypothesized that the response threshold of thermosensory rGCs could be modulated via similar phosphorylation/dephosphorylation cycles upon a temperature upshift.

Five of six residues targeted for phosphorylation in the intracellular domains of GC-A and the sea urchin sperm guanylyl cyclase are conserved in GCY-18 (Figure 3A). We generated a *gcy-18* allele in which all five predicted phosphorylation sites were mutated to alanine (*gcy- 18(5A)*; Figure 3A) via gene editing at the endogenous locus in the *gcy-23 gcy-8* double mutant background. We found that expression of GCY-18(5A) resulted in a significantly lower *T*_Calcium_* both upon short- and long-term temperature upshift (Figure 3B). Examination of the response dynamics in individual neurons showed that following activation at the respective *T*_Calcium_*, intracellular calcium levels appeared to oscillate in a subset of GCY-18(5A)-expressing AFD neurons (Figure 3C). These oscillations were observed primarily upon a rapid temperature upshift but were not observed upon overnight growth at 15°C or 25°C, or in neurons expressing wild-type GCY-18 (Figure 3C). We suggest that phosphorylation of one or more of the targeted residues in GCY-18 may be necessary for both short- and long-term adaptation of the response threshold of this rGC. Moreover, in the absence of this phosphorylation, GCY-18 responses may desensitize rapidly after activation and are re-activated as temperatures rise, resulting in the observed calcium oscillations in response to a rising temperature ramp.

**Figure 3.**
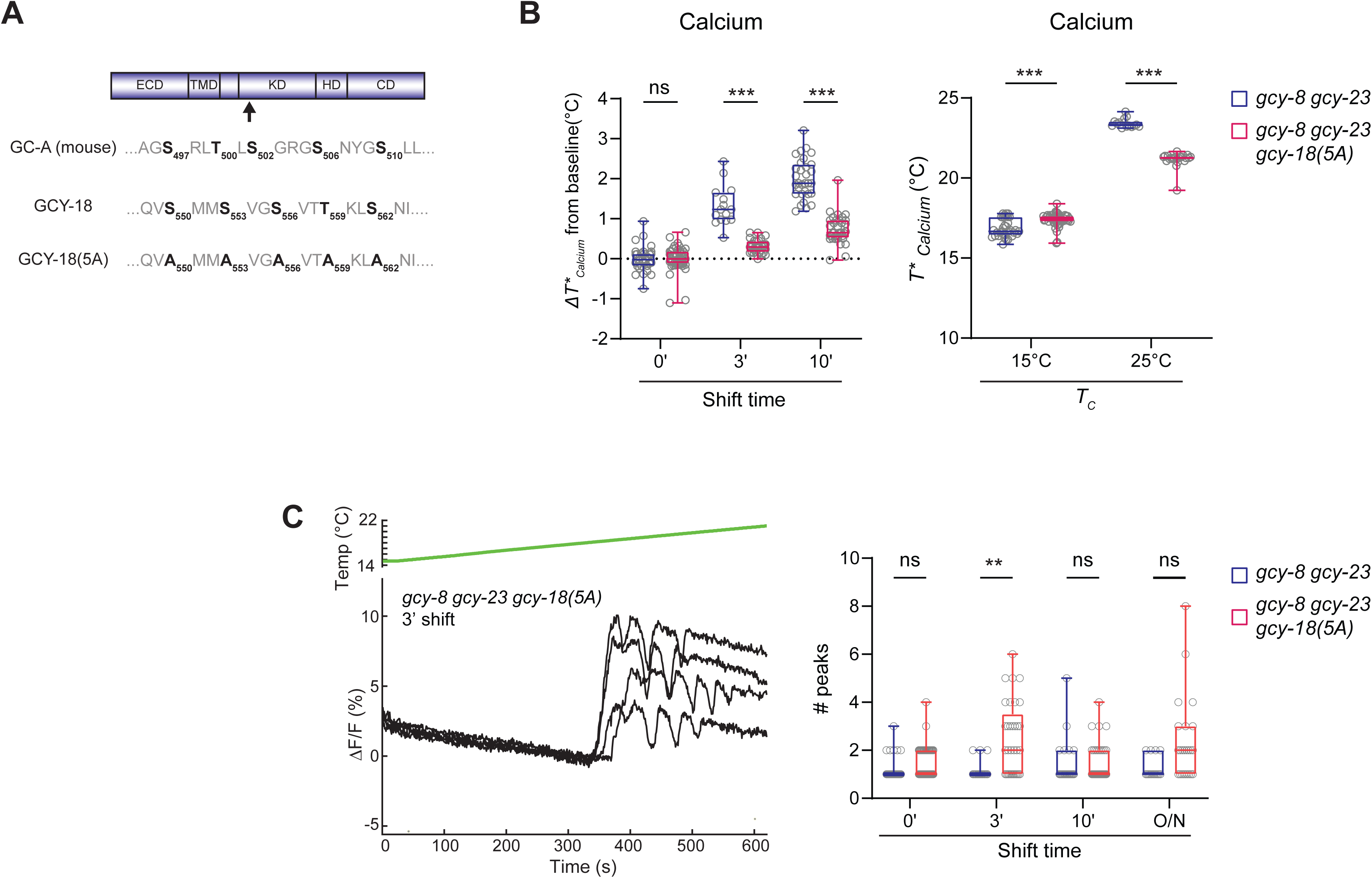
Conserved rGC phosphorylation sites mediate short-term plasticity A) Cartoon illustrating the domain structure of an rGC. The position of conserved phosphorylation sites implicated in desensitization is marked with an arrow. The positions of conserved phosphorylation sites in GC-A, GCY-18, and GCY-18 are marked in bold. Numbers within the sequence mark the position of these sequences in the indicated protein. B) Average changes in *T*_Calcium_* (left) and *T*_Calcium_* (right) in response to a rising temperature ramp in animals of the indicated genotypes subjected to the shown temperature shift conditions. Each circle is the value from a single AFD neuron. C) (Left) Representative traces of GCaMP6s fluorescence changes in individual AFD neurons in *gcy-8 gcy-23 gcy-18(5A)* animals in response to a rising temperature ramp (green; 0.01°C/sec linear ramp) following a 3 min temperature upshift. (Right) Quantification of the number of peaks in fluorescence (see Materials and Methods) in animals of the indicated genotypes subjected to the shown temperature shift conditions. Horizontal lines in boxes in all panels: median, lower and upper quartiles. Whiskers: maximum and minimum values. *, ** and ***: different at p<0.05, 0.01 and 0.001, respectively, between indicated values (t-test). ns – not significant.

### The NCS-2 neuronal calcium sensor may regulate basal cGMP levels in AFD

cGMP and calcium signaling pathways are intricately interconnected in vertebrate phototransduction. Low intracellular calcium levels upon light-induced hyperpolarization activates retinal rGCs via EF-hand containing guanylyl cyclase activating calcium sensor proteins (GCAPs) to regenerate cGMP (39–42). Since buffering of calcium has previously been shown to modulate sensory adaptation dynamics in AFD (17), we asked whether calcium influx feeds back to modulate cGMP in AFD.

Mutations in the *tax-4* CNG channel α subunit largely abrogate temperature-evoked calcium dynamics in AFD (18). We were unable to generate a strain expressing the FlincG3 cGMP sensor in AFD in *tax-4* mutants (also see (26)). Two independent *tax-2* CNG channel β subunit mutants retained partial temperature-evoked calcium responses in AFD although these responses were highly variable (Figure S3A). Temperature-evoked cGMP responses were similarly variable with small amplitudes in *tax-2(p691)* mutants (Figure S3B). Although the response variability and reduced response amplitude precluded accurate measurement of *T*_cGMP_*, these observations suggest that calcium influx may modulate cGMP levels and/or dynamics in AFD.

To determine whether calcium acts via one or more GCAP-like proteins to regulate rGCs in AFD, we next examined the contributions of *C. elegans* neuronal calcium sensor homologs in modulating temperature-evoked cGMP levels in AFD. The *C. elegans* genome encodes three predicted GCAP-related neuronal calcium sensor proteins (NCS-1-3) (43, 44). Transcriptome analyses suggest that NCS-1 and NCS-2, but not NCS-3, are expressed in AFD (45). Although NCS-1 has previously been shown to regulate AFD temperature responses and AFD-driven thermosensory behaviors (27, 46, 47), we observed no defects in calcium or cGMP responses in AFD in *ncs-1* mutants (Figure S3C).

We noted that basal FlincG3 fluorescence levels were significantly higher in *ncs-2* mutants as compared to wild-type animals upon growth at either 15°C or 25°C overnight (Figure 4A). Expression of *gfp* driven under the same promoter driving FlincG3 expression did not result in a similar increase in fluorescence indicating that this increased expression is unlikely to arise from altered expression levels (Figure S3D). While *T*_cGMP_* was not altered in animals under any examined condition (Figure 4B), the amplitude of the cGMP response in AFD in *ncs-2* mutants was increased in animals upon rapid temperature upshift as well as following overnight growth at 25°C (Figure 4C). Increased cGMP response amplitudes and increased basal cGMP levels is predicted to increase the amplitude of the calcium response due to the opening of additional CNG channels, and decrease the threshold of *T*_Calcium_* by enabling the channels to open at a lower temperature following a temperature upshift. As shown in Figure 4D, *T*_Calcium_* adapted to a lower value in two independent *ncs-2* mutants, indicating that within minutes after a temperature upshift, CNG channels open at a lower temperature than in wild-type animals in *ncs-2* mutants. In addition, the amplitude of the calcium response was increased in *ncs-2(ju836)* mutants after 3-10 mins, and in both *ncs-2* alleles upon overnight growth, at 25°C (Figure 4E). We infer that NCS-2 may inhibit basal cGMP levels in both overnight and rapid temperature shift conditions to alter *T*_Calcium_* but not *T*_cGMP_* adaptation.

**Figure 4.**
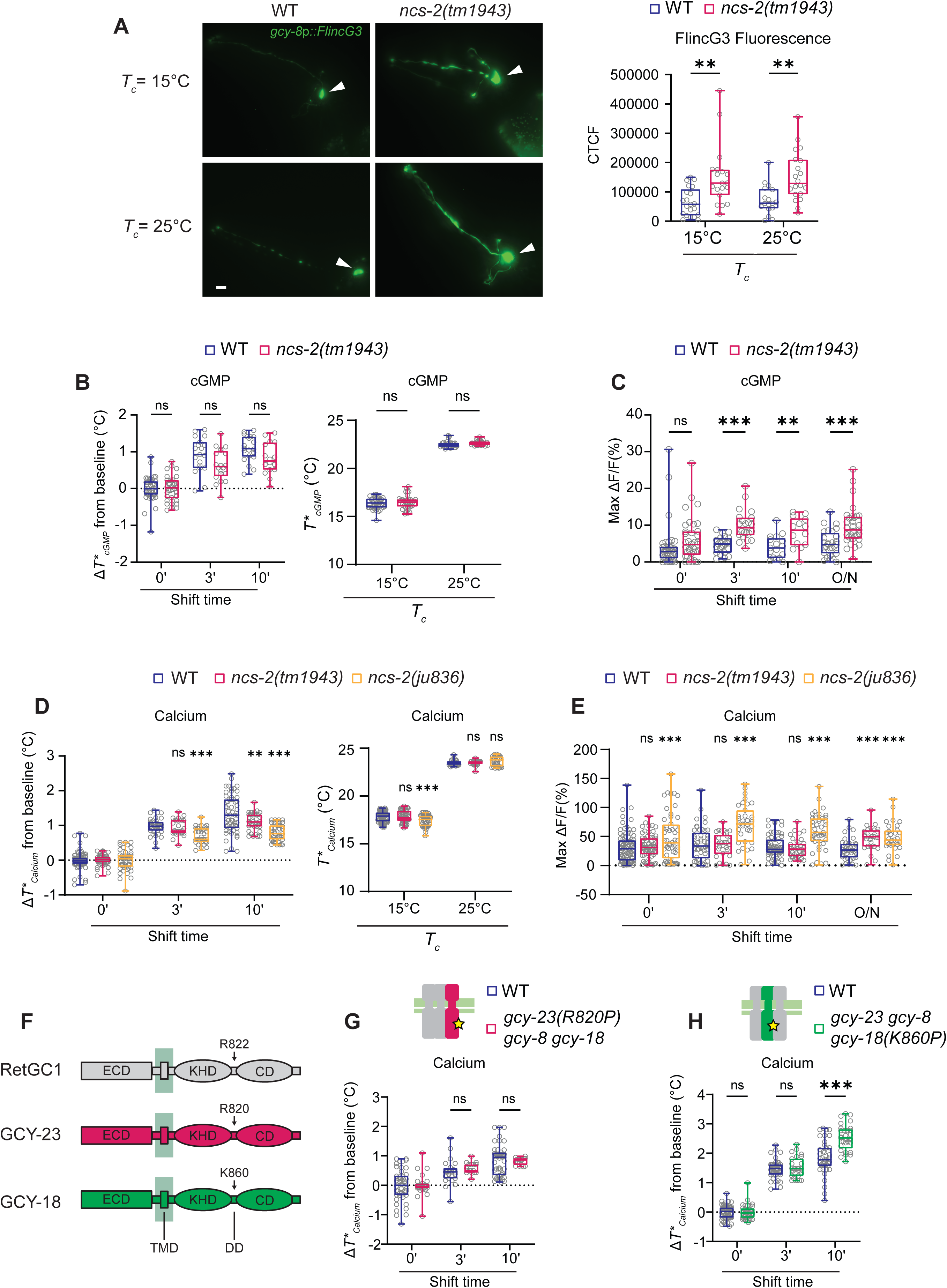
The NCS-2 neuronal calcium sensor protein regulates the catalytic activity of rGCs. **A)** Representative images (left) and quantification (right) of *gcy-8*p::FlincG3 expression in AFD in wild-type and *ncs-2(tm1943)* mutants grown overnight at the indicated temperatures. Arrows indicate AFD soma. Anterior is at top left in all images. Each circle is the fluorescence level in a single AFD. Scale bar: 10 μm. CTCF: corrected total cell fluorescence. **B,D)** Average changes in *T*_cGMP_* (B) and *T*_Calcium_* (D) (left) or *T*_cGMP_* and *T*_Calcium_* values (right) in wild-type or indicated *ncs-2* mutants subjected to the shown temperature shift conditions. Each circle is the value from a single AFD neuron. **(C,E)** Maximum FlincG3 (C) and GCaMP3 (E) response amplitudes in AFD in wild-type and *ncs-2* mutant animals. Each circle is the amplitude of a single AFD neuron. O/N: overnight. **F)** Cartoon of the domains of the indicated proteins and the location of the putative GCAP interacting residues. ECD: extracellular domain, KHD: pseudo-kinase domain, CD: catalytic domain, TMD: transmembrane domain, DD: dimerization domain. Not drawn to scale. **G,H)** Average changes in *T*_Calcium_* in response to a rising temperature ramp in animals of the indicated genotypes subjected to the shown temperature shift conditions. Each circle is the value from a single AFD neuron. Horizontal lines in boxes in all panels: median, lower and upper quartiles. Whiskers: maximum and minimum values. *, ** and ***: different at p<0.05, 00.01, and 0.001, respectively, between indicated conditions or to wild-type (D,E), (t-test except for D,E – ANOVA with Bonferroni correction). ns – not significant.

GCAPs can directly interact with the intracellular domains of retinal rGCs. Although multiple sites of interaction on retinal rGCs have been postulated (48–51) mutating an Arg residue to Pro in the dimerization domain between the kinase homology and catalytic domains of RetGC1 has been shown to abrogate GCAP binding in cultured cells (Figure 4F) (52). We mutated the homologous conserved residues in GCY-18 [GCY-18(K860P)] and GCY-23 [GCY-23(R820P)] (Figure 4F) via gene editing in *gcy-8 gcy-23* and *gcy-8 gcy-18* double mutant backgrounds, respectively, and measured AFD responses. While *T*_Calcium_* in neurons expressing GCY-23(R820P) adapted similarly to neurons expressing wild-type GCY-23 alone (Figure 4G), unlike in *ncs-2* mutants, *T*_Calcium_* adapted to a warmer temperature upon expression of GCY- 18(K860P) (Figure 4H). These observations suggest that NCS-2 is unlikely to interact solely via the targeted residues on either GCY-23 or GCY-18 to modulate their functions, and may act via other as yet unidentified mechanisms to regulate basal cGMP levels in AFD.

### Phosphorylation of the CNG-3 but not TAX-2 channel subunit contributes to rapid thermosensory adaptation

cGMP-mediated opening of CNG channels results in calcium influx; intracellular calcium feeds back via calmodulin to modulate CNG channel properties (53–57). This modulation plays a critical role in regulating adaptation in sensory neurons (12, 58). The TAX-2 and TAX-4 CNG thermotransduction channel subunits have been reported to not contain calcium-calmodulin binding sites (59). However, phosphorylation of TAX-2 by the cGMP-dependent protein kinase (PKG) EGL-4 was shown to be critical for short-term odorant adaptation in the AWC olfactory neurons in *C. elegans* (16). Phosphorylation has also been reported to modulate CNG channel functions in other sensory systems (60–62). We tested whether in addition to adaptation of *T*_cGMP_*, modulation of the response threshold of the CNG channels via PKG-mediated phosphorylation contributes to *T*_Calcium_* adaptation in AFD.

We found that the *tax-2(S727A)* mutation that affects olfactory adaptation only minimally affected *T*_Calcium_* in AFD following a temperature upshift on either a rapid or slow timescale (Figure S4A), suggesting that this residue and/or the TAX-2 subunit are not targets of modulation in AFD. The *C. elegans* genome encodes multiple additional CNG channel subunits in addition to TAX-2 and TAX-4 (59, 63–65). The CNG-3 α subunit is expressed in multiple sensory neuron types including in AFD (64, 65) and has previously been shown to be necessary for robust thermotaxis behavior (66, 67). This channel subunit has also been implicated in the regulation of thermotolerance (64), as well as in short-term olfactory adaptation in the AWC olfactory neurons (59). Temperature-evoked calcium responses in *cng-3(jh113)* null mutants were similar to those in wild-type animals (Figure 5A, 5B), indicating that unlike TAX-2 and TAX-4, this channel subunit is not essential for primary thermotransduction. However, *T*_Calcium_* was consistently lower than in wild-type animals in *cng-3* mutants upon a rapid temperature upshift (Figure 5C). *T*_Calcium_* adaptation was unaltered upon overnight growth at 25°C (Figure 5C). No effects were observed on *T*_cGMP_* adaptation (Figure S4B). The S20 residue in CNG-3 (Figure 5D) has been predicted to be a PKG target mediating short-term olfactory plasticity in the AWC neurons (59). Similar to observations in *cng-3(jh113)* mutants, rapid *T*_Calcium_* adaptation was reduced in animals carrying a S20A mutation in the endogenous *cng-3* locus (Figure 5D; Movie S1), but not upon overnight growth at 25°C (Figure 5D). Consistent with previous observations using *cng-3* null mutants (66, 67), *cng-3(S20A)* mutants also exhibited defects in thermotaxis navigation behaviors such that while wild-type animals grown at 15°C navigated to colder temperatures on a thermal gradient, *cng-3* mutants failed to do so (Figure 5E). These results suggest that phosphorylation of CNG-3 plays a critical role in modulating *T*_Calcium_*, but that this CNG channel subunit is dispensable for primary thermosensory responses.

**Figure 5.**
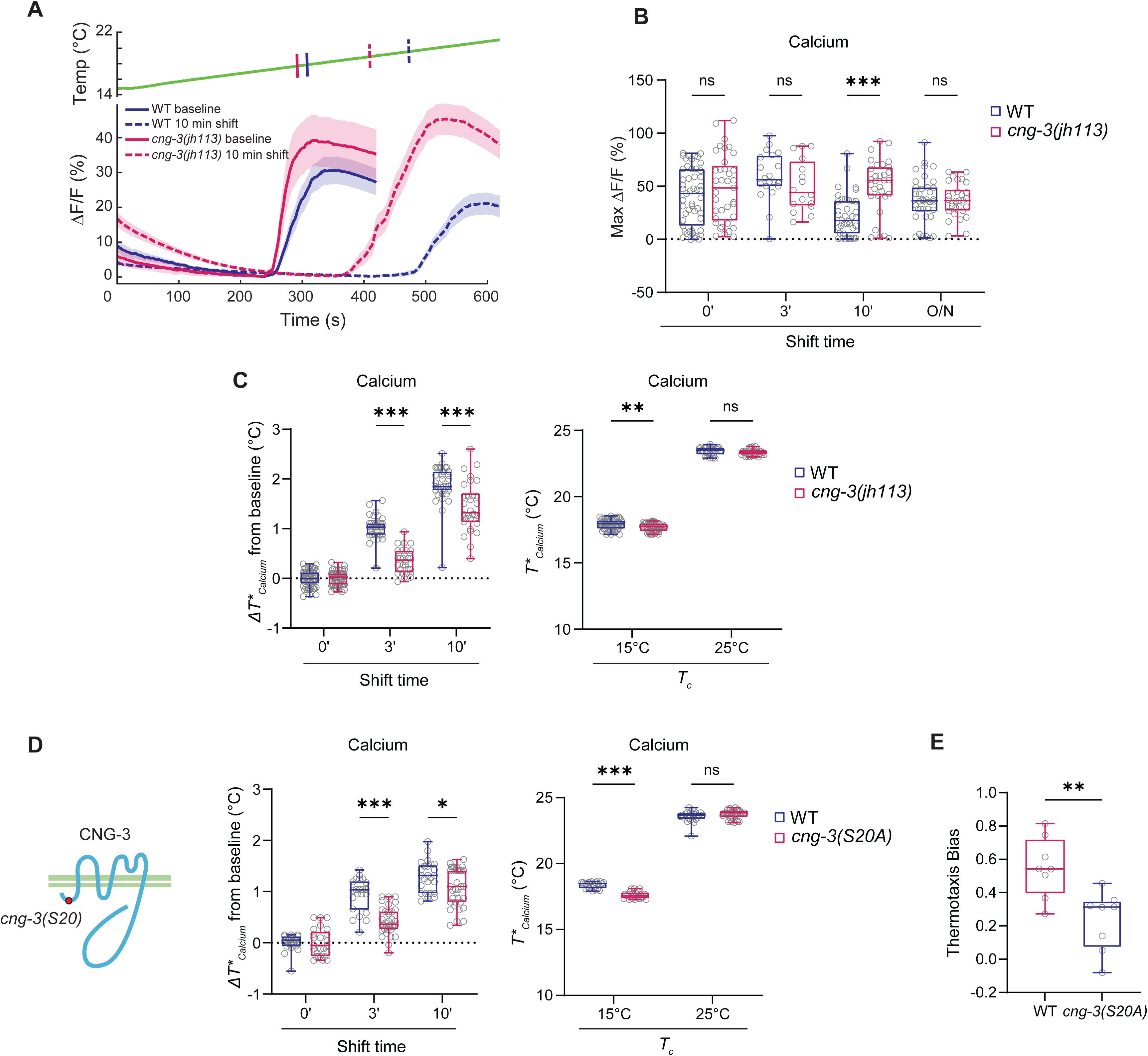
Phosphorylation of the CNG-3 CNG channel subunit is necessary for rapid thermosensory adaptation. **A)** Average changes in GCaMP6s fluorescence in response to a rising temperature ramp (green) in wild-type and *cng-3(jh113)* animals subjected to the shown temperature shift conditions. Shaded regions are SEM. Vertical lines on the temperature trace indicate the quantified average *T*_Calcium_*. **B)** Maximum GCaMP6s response amplitudes in AFD in wild-type and *cng-3(jh113)* mutant animals. Each circle is the amplitude in a single AFD neuron. **C,D)** Average changes in *T*_Calcium_* in response to a rising temperature ramp (left) or average *T*_Calcium_* (right) in wild-type or *cng-3(jh113)* (C), and *cng-3(S20A)* (D) mutants subjected to the shown temperature shift conditions. Each circle is the value from a single AFD neuron. The location of the predicted phosphorylation site in CNG-3 examined here is indicated in the cartoon in (C). **E)** Mean thermotaxis bias of wild-type and *cng-3(S20A)* animals on a thermal gradient. Thermotaxis bias was calculated as (run duration toward colder side – run duration toward warmer side)/total run duration. Animals were cultivated at 15°C and placed on a linear thermal gradient of 19°C-24°C for 30 mins. Each circle represents the thermotaxis bias of a biologically independent assay comprised of 15 animals. Horizontal lines in boxes in all panels: median, lower and upper quartiles. Whiskers: maximum and minimum values. *, ** and ***: different at p<0.05, 0.01, and 0.001, respectively, between indicated conditions (t-test). ns – not significant.

### Two cGMP-dependent protein kinases act redundantly to mediate rapid thermosensory adaptation

The EGL-4 PKG has previously been implicated in mediating multiple forms of olfactory adaptation in AWC in *C. elegans* (4, 16). In particular, EGL-4-mediated phosphorylation of the S727 and S20 residues in TAX-2 and CNG-3, respectively has been suggested to be critical for short-term olfactory adaptation in AWC (16, 59). Given the role of CNG-3 phosphorylation in rapid thermosensory adaptation, we tested whether EGL-4 also regulates *T*_Calcium_* adaptation in AFD.

We found that neither *egl-4(n479 lof)* nor *egl-4(ad450 gof)* mutants exhibited defects in *T*_Calcium_* adaptation upon either a short- or long-term temperature upshift (Figure 6A). In addition to *egl-4*, the *C. elegans* genome encodes a second PKG encoded by *pkg-2* (68). *pkg-2* is poorly characterized and predicted to be expressed in a smaller neuronal subset (45). *pkg-2(tm5814 lof)* animals exhibited relatively minor although significant defects at the 10 min timepoint in *T*_Calcium_* adaptation but not upon overnight growth at any examined temperature (Figure 6B). However, *egl-4; pkg-2* double mutants exhibited strong defects in short- but not long-term *T*_Calcium_* adaptation similar to the phenotypes of *cng-3(jh113)* and *cng-3(S20A)* mutants (Figure 6C). As in the case of *cng-3* mutants, calcium response amplitudes were unaffected in *egl-4; pkg-2* double mutants (Figure S5A). Moreover, *T*_cGMP_* adaptation on either rapid or slow timescales was unaffected in *egl-4; pkg-2* double mutants suggesting that these enzymes do not target the rGCs (Figure 6D). EGL-4 has previously been reported to undergo nuclear translocation only upon prolonged odorant exposure in the AWC olfactory neurons (69). We confirmed that the subcellular localization of a functional GFP-tagged EGL-4 protein was unaltered upon a 15 min temperature upshift in AFD (Figure S5B). These results indicate that the EGL-4 and PKG-2 cGMP-dependent kinases act redundantly to mediate short-term *T*_Calcium_* adaptation possibly via phosphorylation of CNG-3.

**Figure 6.**
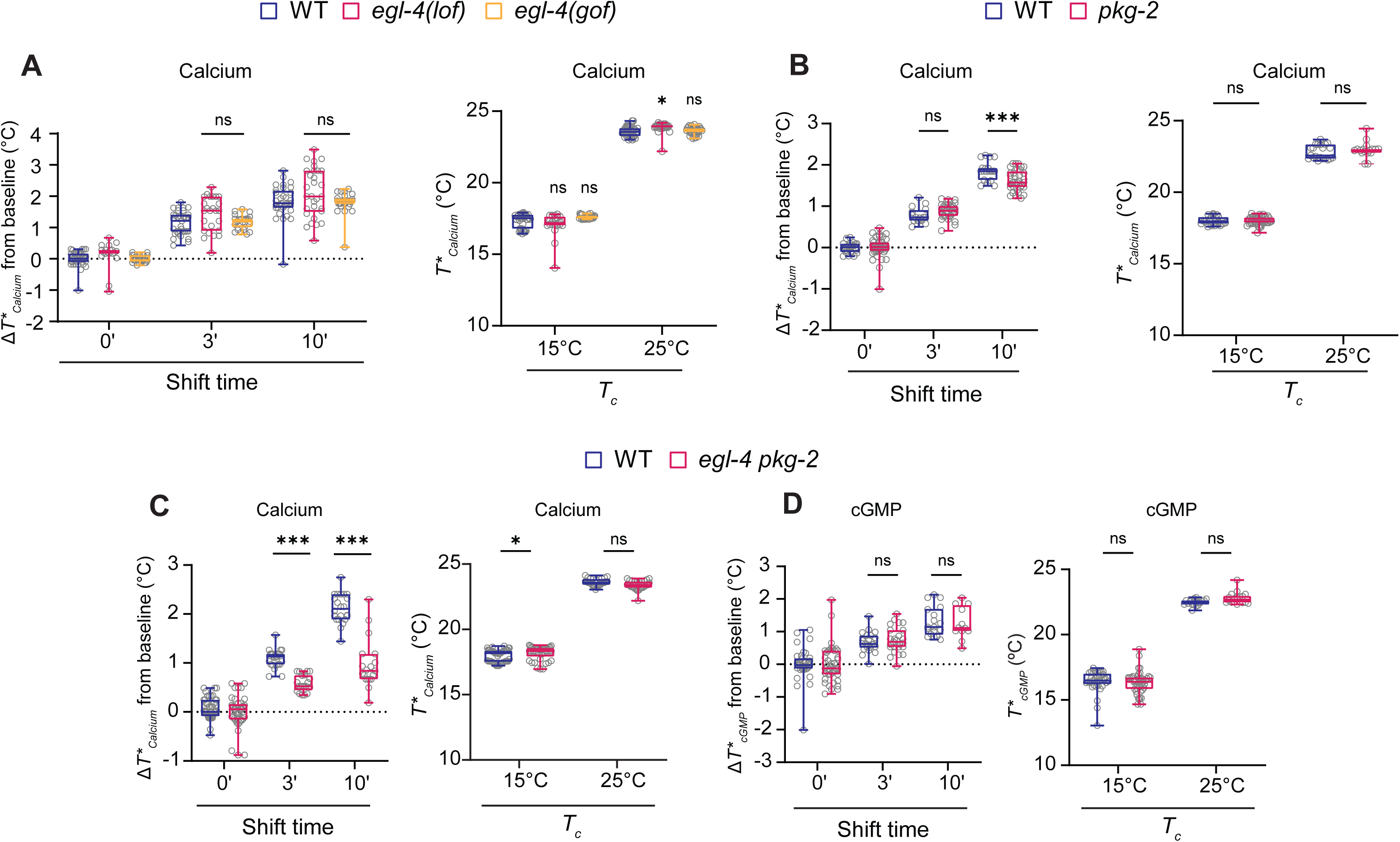
The EGL-4 and PKG-2 PKGs act redundantly to regulate rapid but not long-term thermsosensory adaptation. **A-C)** Average changes in *T*_Calcium_* in response to a rising temperature ramp (left) or *T*_Calcium_* (right) in wild-type or indicated mutants subjected to the shown temperature shift conditions. **D)** Average changes in *T*_cGMP_* in response to a rising temperature ramp (left) or *T*_cGMP_* (right) in wild-type or indicated mutants subjected to the shown temperature shift conditions. Each circle is the value from a single AFD neuron. Alleles used were: *egl-4(n479 lof)*, *egl-4(ad450 gof)*, and *pkg-2(tm5814 lof)*. Horizontal lines in boxes in all panels: median, lower and upper quartiles. Whiskers: maximum and minimum values. * and ***: different at p<0.05 and 0.001, respectively, between indicated conditions or to wild-type (A, left) (t-test in all panels except for A left panel – ANOVA with Bonferroni correction). ns – not significant.

## DISCUSSION

Here we describe the molecular mechanisms that mediate rapid thermosensory response in AFD upon exposure to a temperature upshift (Figure 7). We observe rapid adaptation not only of *T*_Calcium_* as reported previously (10, 17, 20), but also of *T*_cGMP_*, indicating that modulatory mechanisms regulate the first step in the thermosensory signaling pathway to drive response plasticity. We propose that following a temperature upshift, one or more thermosensory rGCs is re-phosphorylated thereby increasing its subsequent response threshold (Figure 7, mechanism 1). cGMP levels feed forward to increase the response threshold of the CNG channels via PKG-mediated phosphorylation of the CNG-3 modulatory subunit (Figure 7, mechanism 2). Calcium influx in turn feeds back through the CNG channels to also regulate rapid adaptation by regulating basal cGMP levels via the NCS-2 neuronal calcium sensor protein (Figure 7, mechanism 3). We suggest that the deployment of multiple mechanisms targeting distinct steps in the thermotransduction pathway allows AFD to rapidly and precisely tune its response properties in response to a brief temperature upshift.

**Figure 7.**
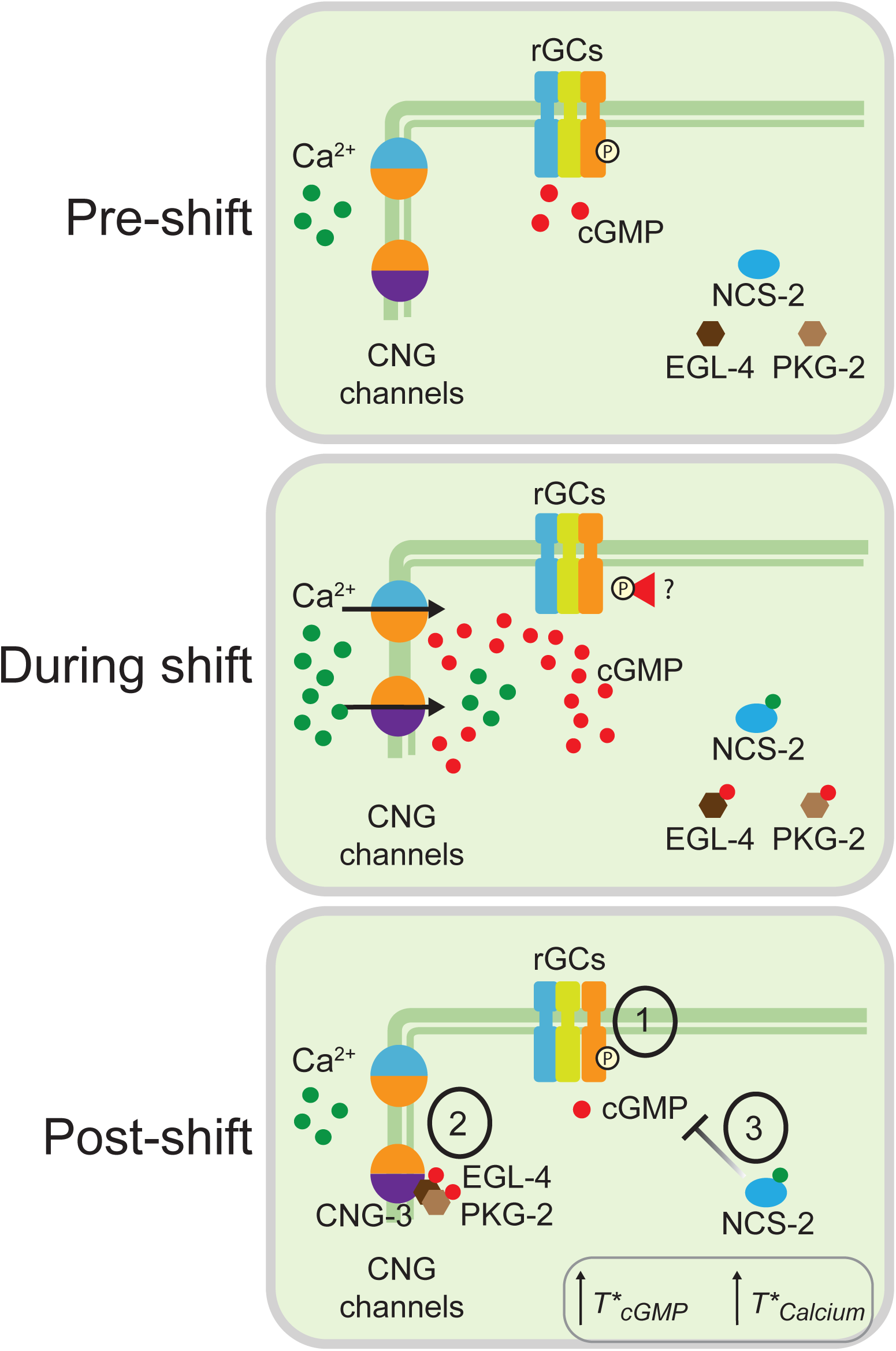
A proposed model of the molecular mechanism of rapid thermosensory adaptation in AFD. Prior to a temperature shift (top), thermosensory rGCs maintain basal levels of intracellular cGMP and the CNG channels are closed. During a temperature upshift (middle), activation of rGCs increases intracellular cGMP levels which open CNG channels to drive calcium influx. rGCs may then be desensitized via dephosphorylation of their intracellular domains via unknown mechanisms. Following the temperature shift (bottom), the threshold of rGC activation may be raised via re-phosphorylation resulting in a higher *T*_Calcium_* and/or *T*_cGMP_* (mechanism 1). cGMP feeds forward to activate the EGL-4 and PKG-2 PKGs which phosphorylate the CNG-3 channel subunit to raise their opening threshold (mechanism 2). Calcium feeds back via NCS-2 to modulate basal cGMP levels and consequently *T*_Calcium_* by targeting rGCs and/or PDEs.

Although only GCY-18 or GCY-23 alone is sufficient to drive rapid adaptation, each rGC drives adaptation at different rates. We find that following a brief temperature upshift, responses in AFD neurons initiate at lower and higher temperatures upon expression of GCY-23- and GCY-18 alone, respectively. The catalytic activity of rGCs is modulated by their phosphorylation state although the required kinases and phosphatases have not been definitively identified (32, 70). Following a temperature upshift, each rGC may be phosphorylated differentially accounting for their characteristic rates of adaptation. The precise adaptation rate of AFD may then be defined via the combinatorial contribution of multiple rGCs. In addition, modulation of the basal catalytic functions of these enzymes via calcium sensor proteins such as NCS-2 may contribute to their ability to drive response plasticity. Thus, rapid shifts in the AFD response threshold are likely underwritten by rapid plasticity in both the rGC response threshold, as well as via modulation of their enzymatic functions to alter CNG channel gating.

While altered *T*_cGMP_* is expected to drive altered *T*_Calcium_*, our results suggest that PKG-mediated phosphorylation of CNG channels specifically contributes to *T*_Calcium_* plasticity. PKG enzymes have been implicated in multiple forms of behavioral plasticity across timescales in different organisms (4, 16, 71–73). Activation of both the EGL-4 and PKG-2 PKGs upon a temperature upshift results in phosphorylation of the CNG-3 but not TAX-2 subunit of the CNG channels to further reset *T*_Calcium_*, likely via reducing the affinity of CNG-3-containing channels for cGMP (59, 61, 62, 74, 75). Loss of function mutations in both *egl-4* and *pkg-2*, as well as mutations in a putative PKG target site in CNG-3, result in similar defects in rapid but not long-term thermosensory adaptation, highlighting the role of this pathway in a temporally defined plasticity paradigm in AFD. The gating, ion conductance, and adaptation properties of CNG channels are regulated by their subunit composition (76, 77), suggesting that modulation of channel subunit composition and properties in a context-specific manner may efficiently drive neuronal plasticity. The positive as well as negative regulation of CNG channels by cGMP in thermosensory adaptation is reminiscent of the incoherent type I feedforward loop network motif that plays a role in generating responses to fold-changes rather than the absolute levels of the stimulus (78, 79), and is distinct from the calcium-mediated negative regulation of CNG channels observed in other sensory systems (6, 12, 58).

Thermosensory adaptation in AFD and odorant adaptation in the AWC olfactory neurons of *C. elegans* share remarkable functional and mechanistic similarities. Both neuron types employ cGMP-mediated signaling pathways to respond sensitively to small stimulus changes over a broad stimulus intensity range although increasing temperature and addition of odorants depolarize and hyperpolarize AFD and AWC respectively (17, 23, 24, 80–82). Similar to the continuous adaptation of the thermosensory response threshold of AFD, the odorant response threshold of AWC also adapts continuously to odor experience on both short and long timescales (4, 10, 11, 16). Although whether odorant adaptation occurs at the level of cGMP production and/or hydrolysis in AWC has not been directly described, PKGs have been implicated in sensory adaptation in both AWC and AFD. While relevant targets necessary for EGL-4-mediated adaptation on a seconds-long timescale are unknown (4), as in AFD, this kinase has been proposed to target CNG subunits including CNG-3 to mediate olfactory adaptation on a timescale of several minutes (4, 16, 59). Moreover, long-term adaptation over hours requires changes in gene expression in both AWC and AFD (10, 28, 29). Nuclear translocation of EGL-4 and the CMK-1 calcium/calmodulin-dependent protein kinase I are necessary for long-term olfactory adaptation and gene expression changes in AWC and AFD, respectively (10, 16, 28, 29, 69, 83). However, in contrast to AWC which has been suggested to only employ calcium-independent feedforward strategies for rapid threshold adaptation (4), AFD appears to employ both feedforward (via cGMP) and feedback (via calcium) adaptation mechanisms. These molecular pathways may represent efficient mechanisms to drive continuous sensory adaptation in neurons utilizing similar sensory transduction machinery.

Results shown here along with previously published work indicate that a partly overlapping set of signaling molecules are targeted to mediate both rapid and slow sensory plasticity in AFD. The functions of both the rGC thermosensors and CNG channels are modulated via posttranslational mechanisms to drive rapid thermosensory adaptation, whereas changes in the expression levels of rGC as well as additional sensory and synaptic genes mediate long-term adaptation (10, 28, 29). While the expression of olfactory receptor genes is not altered by odor history, the expression of CNG channel and calcium pathway genes targeted in rapid adaptation is also regulated by long-term odor experience in mouse olfactory neurons (9). Temporal regulation of functional plasticity in partly overlapping sets of sensory molecules and pathways may allow sensory neurons to precisely adapt their response profiles to the animal’s stimulus history, thereby enabling their remarkable sensitivity and broad dynamic range.

### Limitations of study

Conclusions regarding AFD neuronal activity are limited by using calcium as a proxy for neuronal activity. We are unable to determine absolute calcium or cGMP concentrations in AFD using the FlincG3 or GCaMP6s sensors. Responses to only a 10°C step change from 15°C to 25°C at a single rate of change were examined in this work; distinct pathways may contribute differentially to adaptation in response to other temperature change paradigms. We have not directly measured the cGMP affinities of the channel subunits, and whether this affinity changes upon phosphorylation. Due to the expression of multiple phosphodiesterase subunits in AFD (26, 27, 84), and technical challenges in visualizing cGMP sensor fluorescence in phosphodiesterase mutants (see (26), we have not addressed whether functional changes in phosphodiesterases also contribute to rapid *T*_cGMP_* adaptation.

## MATERIALS and METHODS

### *C. elegans* growth and strain construction

*C. elegans* were maintained using standard conditions on *E. coli* OP50-seeded nematode growth agar plates. The wild-type strain used was *C. elegans* variety Bristol strain N2. The following genetically encoded calcium indicators were used: DCR3055 (*gcy-8*p::GCaMP6s) (20), PY12101 (*gcy-8*p::FlincG3), and PY12108 (*flp-6*p::GCaMP3). PCR-based sequencing was used to confirm the presence of molecular lesions in strains. See Table S1 for the list of all strains used in this work.

### Calcium and cGMP imaging

Calcium imaging was performed essentially as described previously (23, 29). Animals were cultivated at 20°C, and L4 animals were shifted to either 15°C or 25°C overnight. Prior to performing temperature shift experiments, young adult animals grown at 15°C were picked without food onto a 5% agarose pad on a glass coverslip and immobilized with 20mM tetramisole (Sigma Aldrich). A second glass coverslip was placed on top and the sandwich was then placed on a Peltier device on the microscope stage pre-warmed to 25°C. Animals were exposed to 25°C for the desired time period on the microscope stage, following which they were subjected to a temperature ramp from 13°C–26°C at 0.01°C/sec. For 30 min temperature shift experiments, animals were picked onto food-containing growth plates prewarmed at 25°C prior to being moved onto the imaging pads. Calcium imaging data reported in all figures were acquired from AFD soma except in Figure 1 which reports measurements from the AFD sensory endings, and Figure S1C which shows calcium responses from both AFD soma and sensory endings. Typically ∼10 or ∼5 animals were imaged per slide when performing measurements from the soma or sensory endings, respectively.

Current to the Peltier device was controlled through temperature-regulated feedback using a temperature controller (Accuthermo FTC200), an H-bridge amplifier (Accuthermo TX700D), and a thermistor (McShane TR91-170). All videos were acquired using MetaMorph software (Molecular Devices) controlling a Hamamatsu Orca digital camera (Hamamatsu). Imaging was performed using a Zeiss 10X air objective (NA 0.3) on a Zeiss Axioskop2 Plus microscope. ROIs were drawn around the AFD cell bodies or AFD sensory endings. Data were analyzed using custom scripts in MATLAB (Mathworks; https://github.com/SenguptaLab/Hill2023Code). *T*_Calcium_* was defined as the temperature at which ΔF/F increased by a minimum of 2% over at least 8 consecutive seconds with an average slope of >0.3% per second. Analysis of calcium response peaks in *gcy-8 gcy-23 gcy-18(5A)* animals was performed in MATLAB using the *find peaks* function. Traces were put through a lowpass filter with a passband frequency of 0.01 Hz to remove noise before analysis.

Temperature shift experiments prior to cGMP imaging were performed as described for calcium imaging except that animals were subjected to a 13°C-19°C or 13°C-21°C temperature ramp at 0.05°C/sec. All cGMP measurements were performed from the AFD sensory endings. Images were acquired using a Zeiss 40X air objective (NA 0.9). Typically 3-5 animals were imaged per slide and ROIs were drawn around the AFD sensory tip. Data were analyzed using custom scripts in MATLAB (Mathworks; https://github.com/SenguptaLab/Hill2023Code). *T*_cGMP_* was defined as described above for *T*_Calcium_*.

### Fluorescence Imaging

To visualize fluorescence intensities of FlincG3 or *gcy-8*p::GFP, or *gcy-8*p::GFP::EGL-4, animals were cultivated with food at 15°C or 25°C. Animals were then picked onto a 10% agarose pad on a glass slide and immobilized with 20mM tetramisole, and imaged using a 63x oil objective (NA 1.4) on a Zeiss Axio Imager M2 epifluorescent microscope. Fluorescence quantification was performed by measuring the mean fluorescence of an ROI drawn around the AFD soma and subtracting background fluorescence using FIJI in ImageJ. To quantify subcellular localization of GFP::EGL-4 after temperature shift, animals were picked onto food-containing growth plates prewarmed at 25°C prior to being moved onto the imaging pads. Background intensity was subtracted from the mean fluorescence of the nuclear and cytoplasmic regions of the neuron.

### Thermotaxis behavior

Thermotaxis behavior was performed essentially as described (23, 85). Well-fed animals grown overnight at 15°C were transferred to an unseeded NGM plate at 15°C briefly (<10 min) to remove residual bacterial food and then transferred without food onto the assay plate. ∼15 animals were placed at the center of a 10cm unseeded NGM plate pre-equilibrated to the temperature gradient (19°C-24 °C at 0.2°C/cm). Gradients were established by placing assay plates on an aluminum sheet, with temperature maintained using Peltier thermoelectric temperature controllers (Oven Industries). The temperature at both ends of the gradient was measured using a two-probe digital thermometer (Fluke Electronics) at the beginning of each imaging session. Animal movement was recorded for 35 minutes at 1Hz using a PixeLink CCD camera controlled by custom MATLAB scripts. The first 5 minutes of recording were discarded, and videos were analyzed using custom MATLAB scripts. Analysis code can be found at https://github.com/SenguptaLab/Hill2023Code.

### CRISPR/Cas9-mediated gene editing

All cRNAs, tracrRNAs and Cas9 protein were obtained from Integrated DNA Technologies (IDT). Point mutations in *gcy-18*, *gcy-23*, *tax-*2, and *cng-3* were generated using ssODN repair templates which included the mutated site along with 35 bp 5′ and 3′ homology arms. Injection mixes for all manipulations included the ssODN repair template (110 ng/μl), Cas9 protein (250 ng/μl), tracerRNA(100 ng/μl), crRNA(28 ng/μl) and a co-injection marker (*myo-3*p::mCherry (50ng/μl)). F1 animals expressing the co-injection marker were analyzed for the presence of the desired mutations via PCR and sequencing of the relevant genes. F2 progeny were subsequently screened for homozygous edits via PCR and sequencing. Sequences of all oligonucleotides are listed in Table S2.

### Molecular biology

The *gcy-8*p*::gfp::egl-4* plasmid was constructed by replacing the promoter in the *odr-3*p*::gfp::egl-4* plasmid (Addgene) with the AFD-specific *gcy-8* promoter using standard restriction enzyme-mediated cloning, and verified by sequencing.

### Statistical analyses

All statistical analyses were performed using Graphpad Prism version 10.0.0 (www.graphpad.com) which was also used to generate all box plots. Statistical test details and the number of analyzed samples are reported in each figure legend.

## Acknowledgments

We thank the *Caenorhabditis* Genetics Center and the National BioResource Project (NBRP) for strains. We are grateful to the Sengupta lab for experimental and conceptual advice and input, Charmi Porwal for technical assistance, and Nathan Harris, Sam Bates, and Alison Philbrook for critical comments on the manuscript. We acknowledge receipt of reagents from Daniel Colón-Ramos and Noelle L’Etoile. This work was funded in part by the NIH (R35 GM122463 – P.S.).

## SUPPLEMENTAL FIGURE LEGENDS

**Figure S1.**
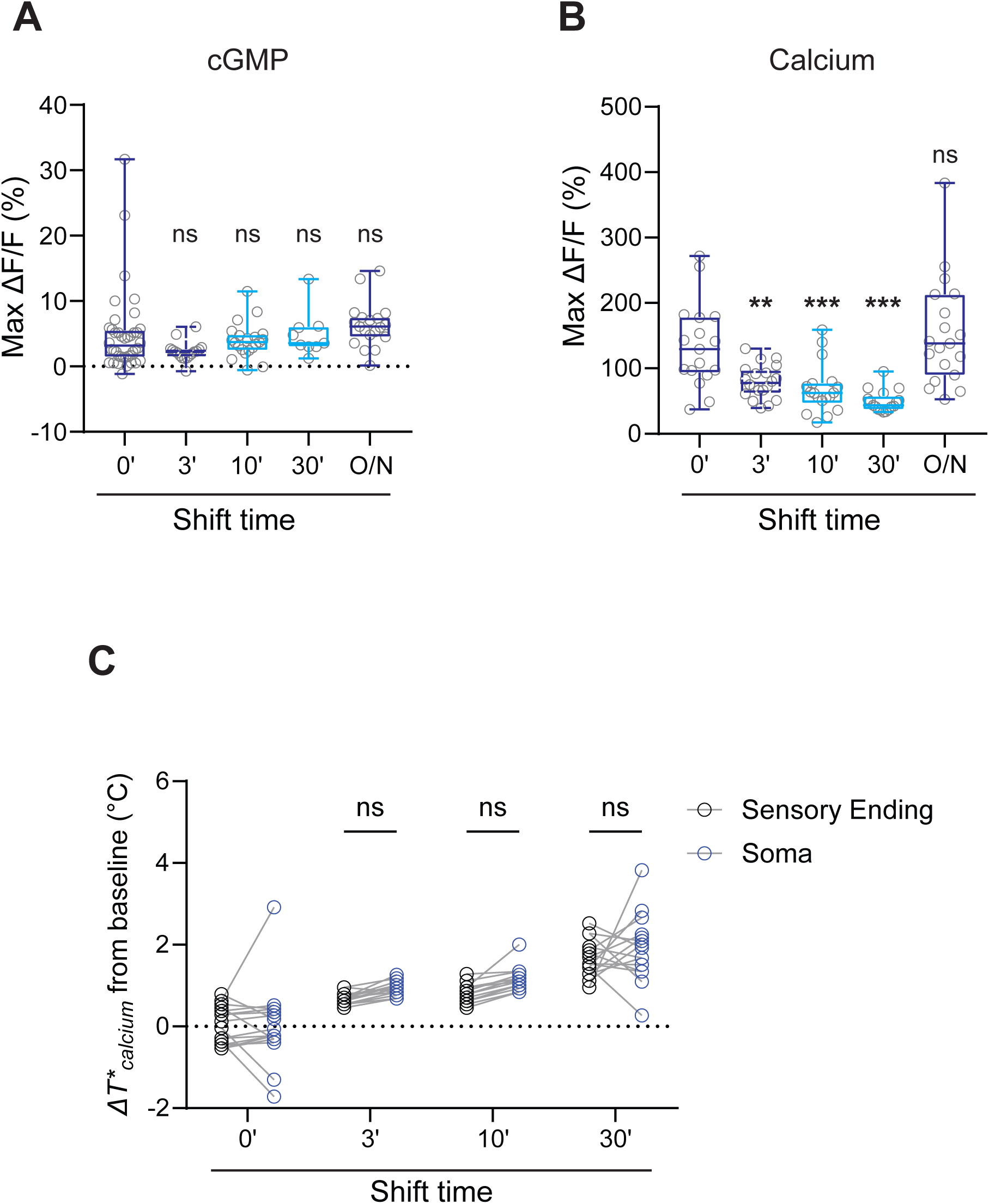
Response amplitudes of temperature-evoked cGMP and calcium dynamics in AFD upon a temperature upshift. **A,B)** Maximum amplitudes of cGMP (A) and calcium (B) responses in AFD in the shown temperature conditions (from average traces shown in Figure 1C-D). Each circle is the maximum response amplitude of a single AFD neuron. Horizontal lines in boxes: median, lower and upper quartiles. Whiskers: maximum and minimum values. ***: different at p<0.001 from the 0’ condition (ANOVA with Bonferroni correction). ns – not significant. O/N – overnight. **C)** Average change in *T*_Calcium_* from the baseline *T\** upon a temperature upshift for the indicated time periods. Measurements were performed from the sensory endings or cell bodies of AFD in each individual animal. ns – not significant.

**Figure S2.**
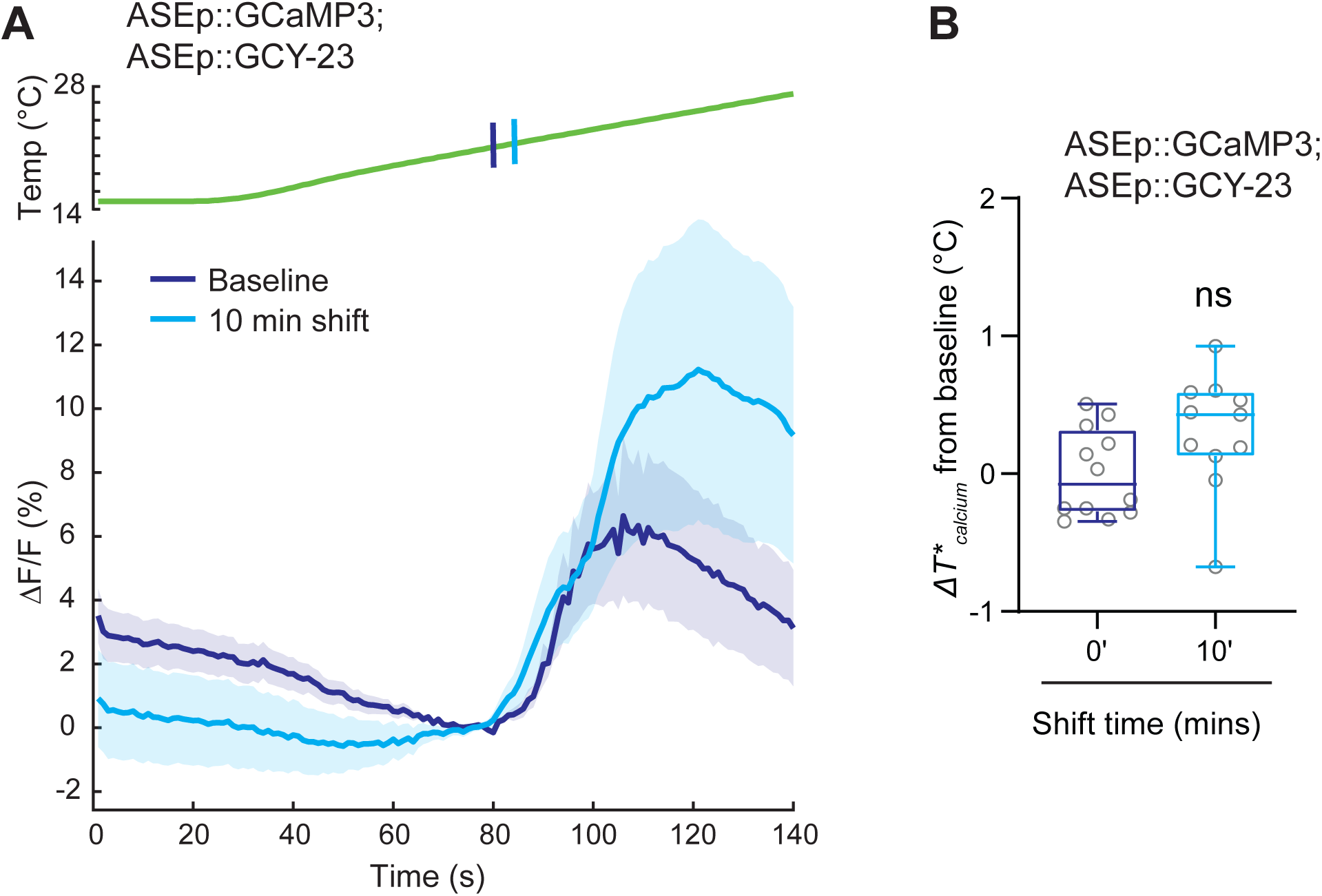
Misexpression of GCY-23 does not confer rapid temperature adaptation. (Left) Average changes in GCaMP3 fluorescence in ASE chemosensory neurons misexpressing *gcy-23* in response to a rising temperature ramp (green; 0.05°C/sec linear ramp). Animals were grown overnight at 15°C or shifted to 25°C for 10 mins prior to imaging. Shaded regions are SEM. Vertical lines on the temperature trace indicate the quantified average *T*_Calcium_*. *gcy-23* was expressed in ASE under the *flp-6* promoter (23). (Right) *T*_Calcium_* of *flp-6*p*::gcy-23* expressing animals grown overnight at 15°C or shifted to 25°C for 10 mins. Each gray circle is the value in a single neuron. Horizontal lines in boxes: median, lower and upper quartiles. Whiskers: maximum and minimum values. ns – not significant (t-test).

**Figure S3.**
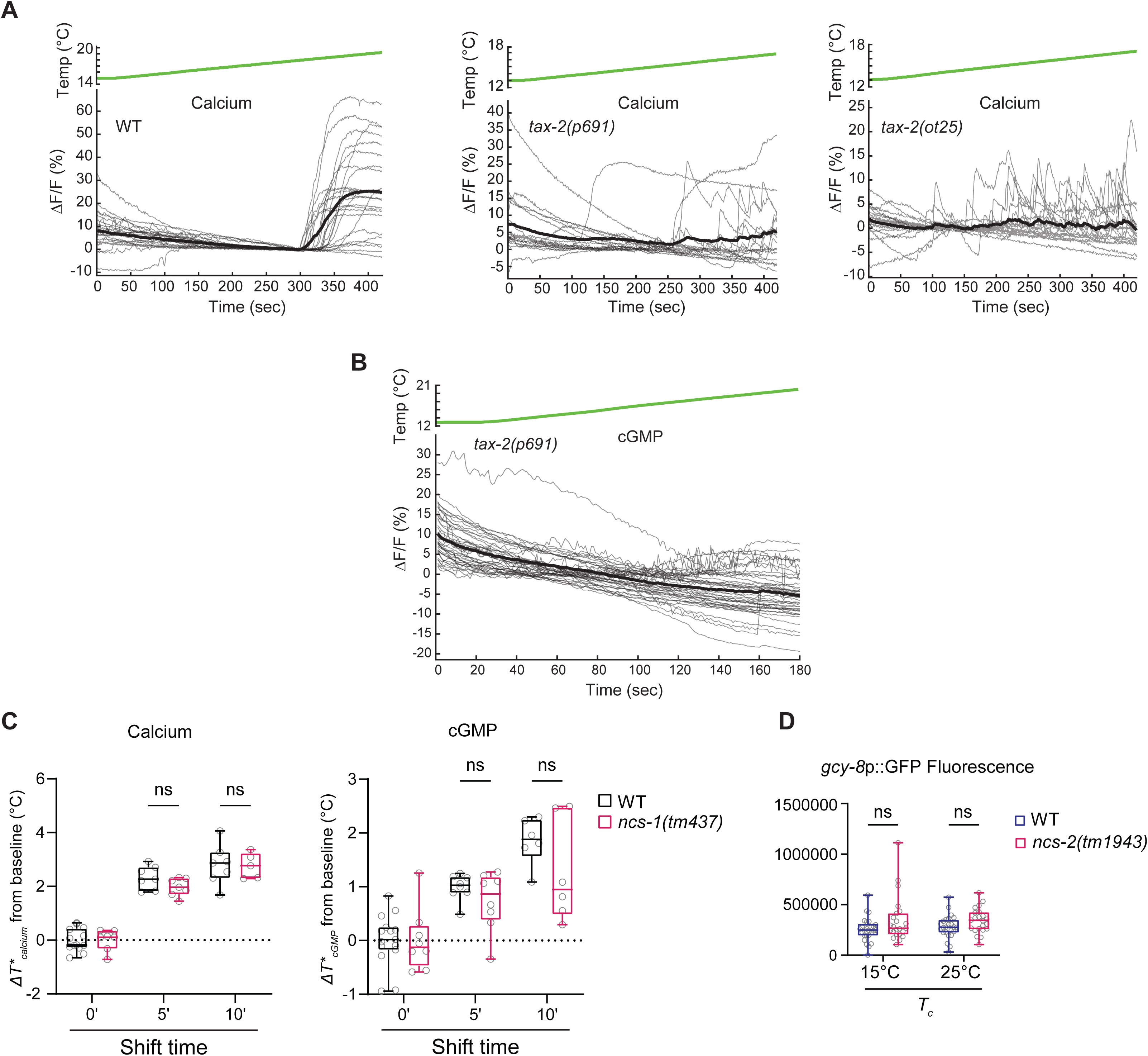
Temperature responses of *tax-2* and *ncs-1* mutants. **A,B)** Average changes in GCaMP6s (A) or FlincG3 (B) fluorescence in response to a rising temperature ramp (green; 0.05°C/sec linear ramp) in animals of the indicated genotypes grown overnight at 15°C. Thick and thin lines are the average and individual fluorescence responses, respectively. Wild-type responses in A are repeated from Figure 2J. **C)** Average changes in *T*_Calcium_* (left) or *T*_cGMP_* (right) upon a temperature upshift for the indicated time periods in wild-type or *ncs-1* mutants. Each circle is the value from a single AFD neuron. Horizontal lines in boxes: median, lower and upper quartiles. Whiskers: maximum and minimum values. ns – not significant. **D)** Quantification of *gcy-8*p::GFP expression in AFD in wild-type and *ncs-2(tm1943)* mutants grown overnight at the indicated temperatures. Each circle is the fluorescence level in a single AFD. CTCF: corrected total cell fluorescence.

**Figure S4.**
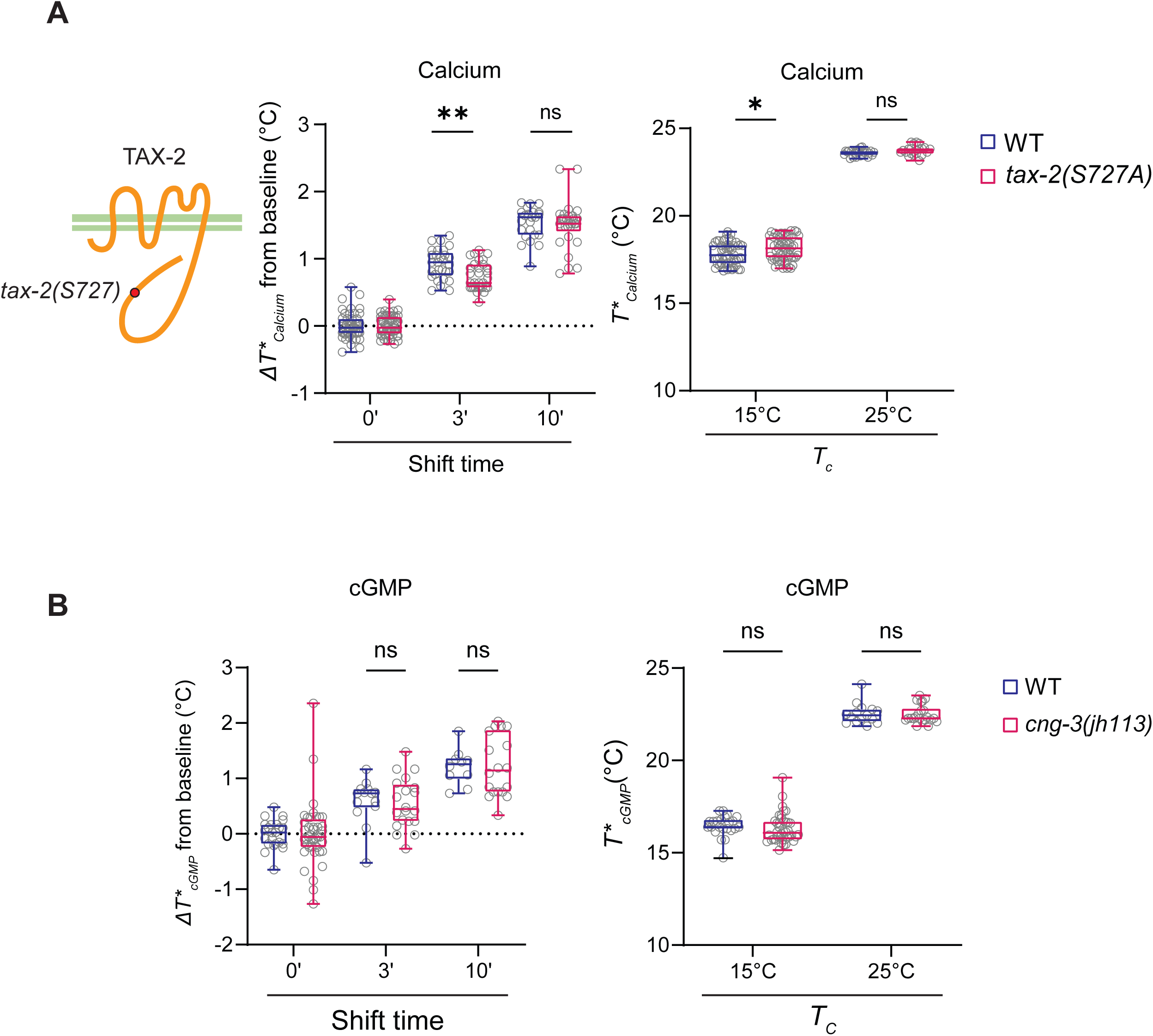
Rapid adaptation is unaffected in *tax-2* phosphomutants. **A)** (Left) Cartoon of TAX-2 showing the locations of the predicted phosphorylation site examined here. (Right) Average changes in *T*_Calcium_* in response to a rising temperature ramp or average *T*_Calcium_* in wild-type or *tax-2(S27A)* mutants subjected to the shown temperature shift conditions. Each circle is the value from a single AFD neuron. **B)** Average change in *T*_cGMP_* from the baseline *T\** (left) or *T*_cGMP_* in wild-type and *cng-3(jh113)* null mutants subjected to the shown temperature shift conditions. Each circle is the value from a single AFD neuron. Horizontal lines in boxes in all panels: median, lower and upper quartiles. Whiskers: maximum and minimum values. * and **: different at p<0.05 and 0.01, respectively, between indicated conditions (t-test). ns – not significant.

**Figure S5.**
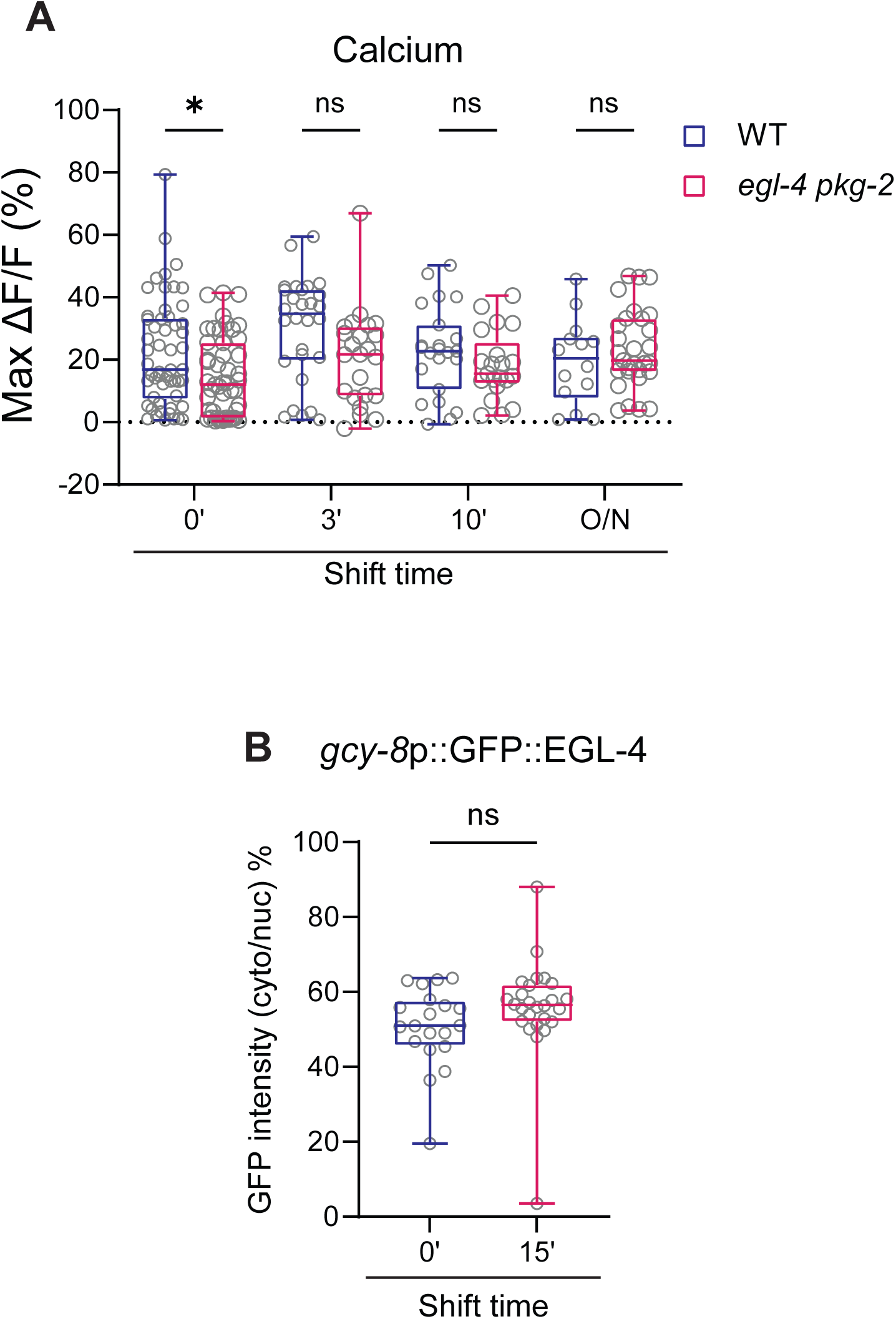
Calcium responses in *egl-4 pkg-2* double mutants. **A)** Maximum GCaMP3 response amplitudes in AFD in wild-type and *egl-4 pkg-2* double mutant animals. Each circle is the amplitude of a single AFD neuron. Horizontal lines in boxes: median, lower and upper quartiles. Whiskers: maximum and minimum values. *: different at p<0.05 between indicated conditions (t-test). ns – not significant. **B)** Subcellular localization of GFP::EGL-4 expressed in AFD. Each circle is the percentage of cytoplasmic to nuclear fluorescence in a single AFD neuron.

**Movie S1**. Calcium responses in two wild-type (bottom) and two *cng-3(S20A)* mutants (top) expressing GCaMP6s separated by a non-transgenic animal. The temperature ramp is indicated at top left. Anterior is at left. Images were captured at 1 Hz and played back at 30 frames per sec.

**Table S1.**
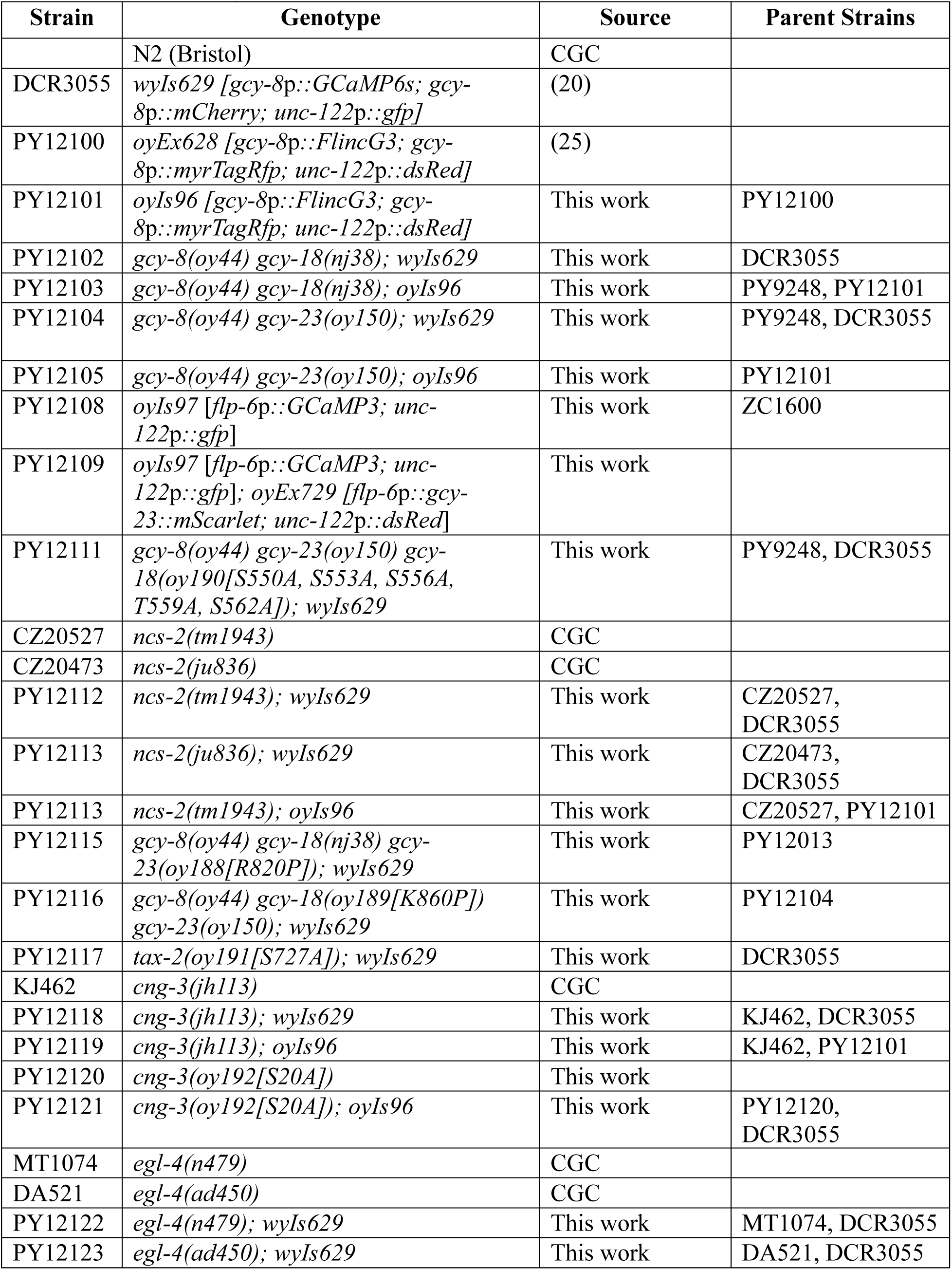
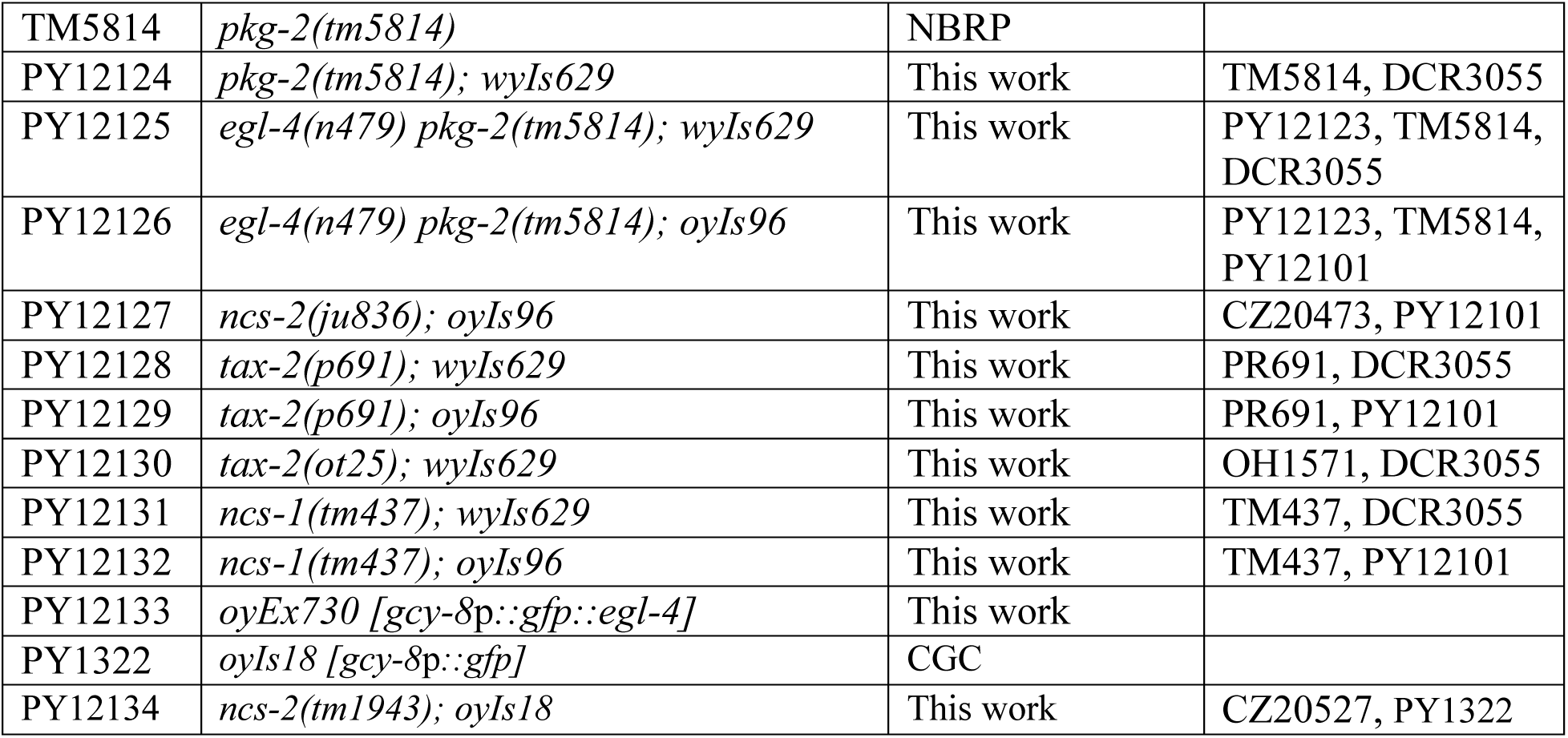
List of *C. elegans* strains used in this work.

**Table S2.**
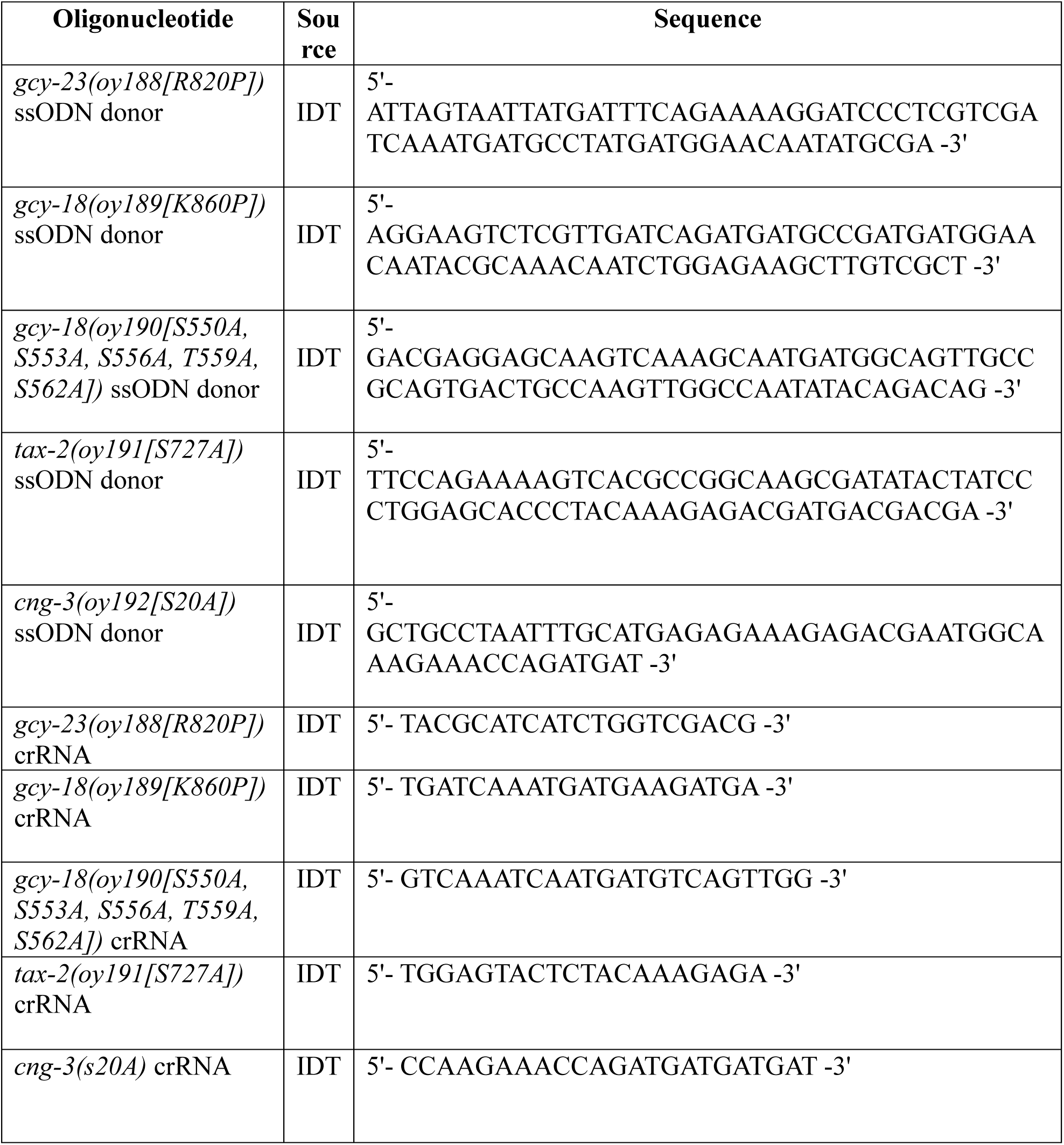
List of primers used in this work.

